# A theoretical model of mitochondrial ATP synthase deficiencies. The role of mitochondrial carriers

**DOI:** 10.1101/2021.07.13.452228

**Authors:** Jean-Pierre Mazat, Anne Devin, Edgar Yoboue, Stéphane Ransac

## Abstract

The m.8993T*>*G mutation of the mitochondrial *MT-ATP6* gene is associated with NARP syndrome (Neuropathy, Ataxia and Retinitis Pigmentosa). The equivalent point mutation introduced in yeast *Saccharomyces cerevisiae* mitochondrial DNA considerably reduced the activity of ATP synthase and of cytochrome-c-oxidase preventing yeast growth on oxidative substrates. The over-expression of the mitochondrial oxodicarboxylate carrier (Odc1p) is able to rescue the growth on oxidative substrate by increasing the substrate-level phosphorylation of ADP coupled to conversion of *α*-ketoglutarate (AKG) into succinate with an increase in Complex IV activity. Previous studies showed that equivalent point mutations in ATP synthase behave similarly and can be rescued by Odc1p overexpression and/or uncoupler of OXPHOS from ATP synthesis. In order to better understand the mechanism of ATP synthase mutation bypass, we developed a core model of mitochondrial metabolism based on AKG as respiratory substrate. We describe the different possible metabolite output and the ATP/O ratio values as a function of ATP synthase inhibition.

## 1. Introduction

Mitochondria support aerobic respiration and produce most of cellular ATP by oxidative phosphorylation, i.e. the coupling of a series of redox reactions, the electron transport chain (ETC) with the ATP synthase through a transmembraneous proton gradient. The ETC is mainly fed by the NADH and succinate generated in the TCA cycle and some other possible dehydrogenases.

The ATP synthase organizes into a matrix domain (F1 inside mitochondria) where ATP is synthesized and a membrane-embedded domain (Fo) that moves protons across the membrane **[1–4]**. Dozens of point mutations in the mitochondrial MT-ATP6 gene (coded by ATP6 gene in yeast mtDNA) have been identified as leading to deleterious neuromuscular disorders **[5]**. This gene codes for the subunit a in the ATP synthase. It is in contact with the c-ring in the Fo membrane domain and has been proposed to optimize the transmembrane conduction of protons **[6,7]**. The m.8993T>G mutation of the mitochondrial MT-ATP6 gene affecting mitochondrial energy transduction has been particularly studied **[8]**. This mutation has been associated in human with numerous cases of neuropathy, ataxia and retinitis pigmentosa (NARP) and maternally inherited Leigh syndrome.

The equivalent mutations introduced in the yeast *Saccharomyces cerevisiae* (NARP mutant) **[9]** dramatically slow down its respiratory growth. It was recently shown that this mutation prevents proton release in the mitochondrial matrix strongly decreasing ATP synthesis to 10% of the wild type (WT) value. **[6]**. Concomitantly with the ATP synthase defect, a decrease in Complex IV amount is observed (retrograde signaling pathway linking ATP synthase and complex IV biogenesis **[8,10,11]**).

An additional molecule of ATP (in yeast) or GTP (in liver) is produced in TCA cycle by Succinyl-CoA synthetase, also named succinate thiokinase. It is called substrate level phosphorylation (SLP) because it involves the transfer of a phosphate from a donor molecule (here the succinyl-Pi) to ADP to form ATP. Following the pioneer work of Schwimmer et al. **[10]**, a rescue of the NARP mutation in yeast was mediated through the mitochondrial carrier Odc1p overexpression **[8]**. Odc1p is a mitochondrial carrier that exchanges 2-oxoglutarate (AKG) against succinate, malate or citrate (and several other metabolites, not *a priori* involved in the rescuing process) **[12]**. Odc1p overexpression presumably increases AKG uptake in mitochondria promoting the subsequent increase in ATP synthesis by SLP (SLP-ATP) **[10]**. In both cases the rescue is accompanied by an increase of complex IV amount without full restoration of the WT value. A model of the interdependence between ATP synthase and complex IV amount via the increase in mitochondrial membrane potential has been proposed in **[8]** in relationship with mitochondrial membrane hyperpolarization (see also **[13]**).

Odc1p carrier was identified and characterized by Palmieri et al. **[12]**. It operates by a strict counter exchange mechanism. Odc1p carrier can principally uptake 2-oxoglutarate against itself (112.5), succinate (19), malate (56) and citrate (28.6). The numbers in brackets indicate in mmol/min/g protein the rate of 2-oxoglutarate uptake in proteoliposomes preloaded with the corresponding metabolite (table II in **[12]**). The diversity of output catalyzed by Odc1p offers different metabolic solutions to enhance SLP-ATP. In order to have a better understanding of the rescue mechanism by Odc1p we developped in this paper a metabolic model of yeast mitochondria that characterizes the different possible metabolic pathways bypassing oxidative phosphorylation ATP deficiency in isolated yeast mitochondria using α-ketoglutarate (AKG) as respiratory substrate. We show that several metabolite outputs following AKG input are possible and depend upon the activity of the ATP synthase. We also show that the simple property of Odc1p as a transporter is not sufficient to explain the ATP synthase rescue. We propose some new hypotheses to explore. Finally we discuss the therapeutic application of this study in human.

## 2. Materials and Methods

The model of mitochondrial metabolism was written with Copasi [14] and is detailed in supplementary materials S1. It involves a relevant representation of the coupling between respiratory chain and ATP synthesis through a chemiosmotic proton gradient ΔµH^+^ considered here as a pseudo substrate called PMF (for Proton Motive Force). For those reactions which depend only on the ΔpH or on the Δψ component of the ΔµH^+^, we take, following Bohnensack [15], ΔpH = 0.2 PMF and Δψ = 0.8 PMF. In our model, the PMF, as postulated by Mitchell’s chemiosmotic theory [16], is the result at steady-state of the opposite activities of the respiratory chain on the one hand and of the ATP synthase on the other hand even though some recent results indicate a more complex mechanism [17]. Copasi also allows the calculation of EFMs (Elementary Flux Modes, i.e. the minimal set of fluxes at steady state) [18,19] but with integer stoichiometric coefficients. This explains why some reactions have high stoichiometric coefficients and why some coefficients of the reactions in the EFMs are high (supplementary material S2). The kinetic parameters of the rate equations are listed in Supplementary Materials S1.

The model includes the different exchanges catalyzed by Odc1p under the form of 3 reactions for AKG input:

**T2**: AKGc + MALm = AKGm + MALc.

**T22**: AKGc + SUCCm = AKGm + SUCCc.

**T23**: AKGc + CITm = AKGm + CITc.

In addition the dicarboxylate carrier (DIC) catalyzes the exchanges:

**T31:** MALm + Pic = MALc + Pim.

**T33:** SUCCm + Pic = SUCCc + Pim.

## 3. Results

The aim of the model was to simulate the experiments on isolated yeast mitochondria with 10 mM AKG as respiratory substrate and to characterize all possible metabolic pathways inside mitochondria, and metabolite outputs including ATP synthesis. We also studied the oxygen consumption flux and calculated the ATP/O ratio for which experimental results already exist [20].

In a first part we will describe the Elementary Flux Modes (EFMs) i.e. the minimal pathways at steady-state inside a metabolic network [18]. In this context, minimal means that the removal of a step prevents the establishment of a steady state. They either connect input with output metabolites or are cycles at steady-state. Any set of fluxes at steady-state in a metabolic network can be written as a non-negative linear combination of EFMs [18,21].

### 3.1. Determination of EFMs with 2-Oxoglutarate (AKG) as respiratory substrates

The EFMs are described in Supplementary Materials S2. The direction of the carriers in the model are such that only AKG is allowed as respiratory substrate. In these conditions, we obtain 15 possible EFMs. Among these 15 EFMs, 8 involve the input of only AKG, ADP, Pi and O, and the output of ATP (represented in Fig. 1 and in supplementary Materials S2). The outputs are succinate only (EFMs 3 and 11, Fig. 1 a and d respectively), malate only (EFMs 4 and 12, Fig. 1b and e respectively), malate + citrate (EFMs 6 and 13, Fig. 1c and f), Succinate + citrate (EFM 14, Fig. 1g) or succinate + malate + citrate (15, Fig. 1i). EFMs 11, 12 and 13 involve a membrane leak (L) which consumes the PMF without ATP synthesis (ASYNT = 0) with a rather low ATP/O of 0.5 or 1. There is a last EFM without oxygen consumption, EFM 5 represented in Fig. 1h. This EFM involves the reversion of ATP synthase (-ASYNT). Even though an oxygen consumption is experimentally observed, this EFM cannot be excluded because it can participate to a linear combination of EFMs, with a net consumption of oxygen. All in all the ATP/O values range from 0.5 to 6. Two EFMs (EFM 3 and EFM 6) display an ATP/O of 2.15 close to the published value of 2.3 [10,20]. However because the actual fluxes in a metabolic network can be seen as a linear combination of EFMs, the experimental value of 2.3 can result from a combination of different EFMs with different ATP/O values. Thus, any of the 9 EFMs can be implicated either alone (EFMs 3 or 6, figure 1a or c) or in combination.

**Figure 1.**
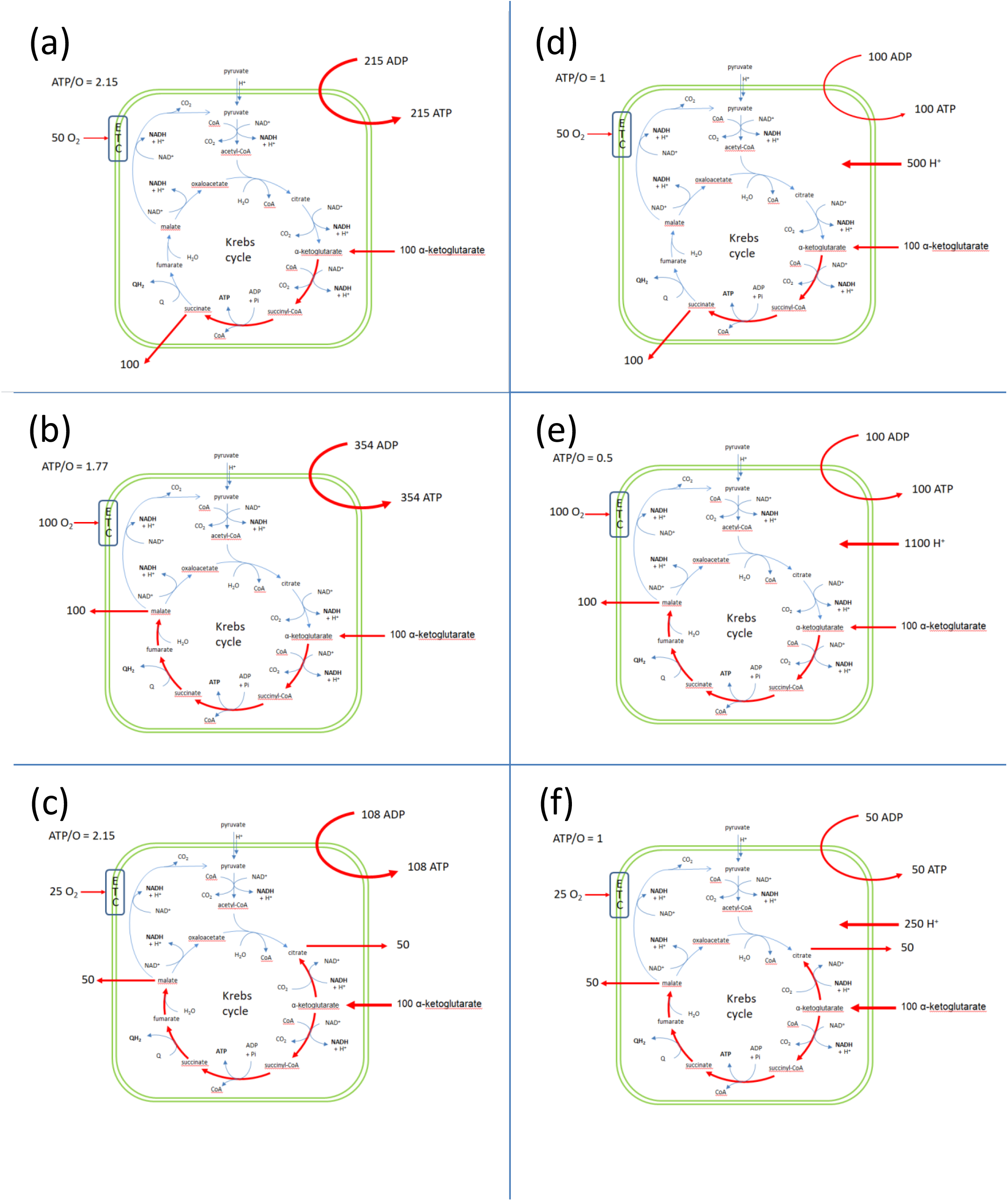
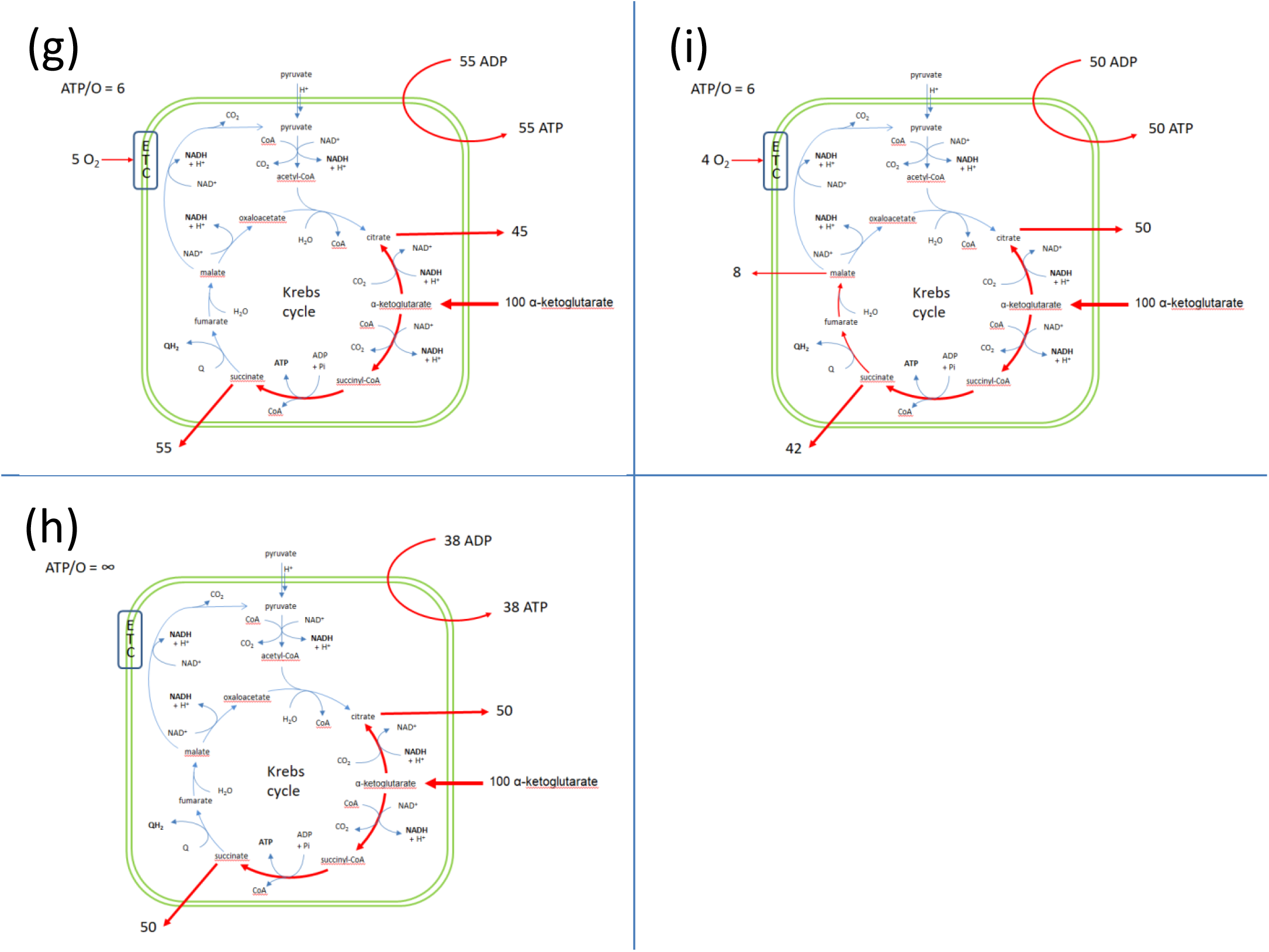
Schematic representation of the EFMs that explain the set of fluxes in isolated yeast mitochondria respiring on AKG. (d), (e) and (f) correspond to a consumption of PMF through the leak L. All values are given for an input of 100 AKG per time unit (in arbitrary units) in order to facilitate the comparison of the release fluxes. The EFMs are detailed in Supplementary Material S2.

To get more insight in the actual metabolic fluxes and the pathway taken by AKG inside the mitochondria, we developped a kinetic model using Copasi [14] involving the kinetic properties known for each step as described in supplementary material S1.

### 3.2. Dynamics of oxygen consumption and ATP generation and determination of the ratio ATP/O as a function of ATPsynthase activity (Fig. 2)

Three properties of the model must be emphasized.

1. Because the thermodynamic span between the pair NADH/NAD and QH_2_/Q is much greater than the span between succinate/fumarate and QH_2_/Q, NADH is oxidized before QH_2_ (Fig. 2b) and the accumulation of QH_2_ blocks the respiratory complex II (RCII or SDH).
2. When the ratio NADH/NAD is high (low ATP Synthase activity) the reversion of IDH3 associated with NADH reoxidation is possible.
3. When both NADH/NAD and QH_2_/Q ratios are low (high activity of the respiratory chain and ATP synthase), RCII is active and succinate is easily transformed in fumarate competing with the output of succinate against AKG through Odc1p. The AKG/succinate exchange is thus decreased and largely replaced by the AKG/malate exchange (Fig. 2a at high ASYNT activity).

Let us now describe the behavior of mitochondrial metabolism with AKG as respiratory substrate as a function of ATP synthase activity (corresponding to different degrees of activity or different amount of the ATP synthase inhibitor oligomycin) (Fig. 2).

**Figure 2.**
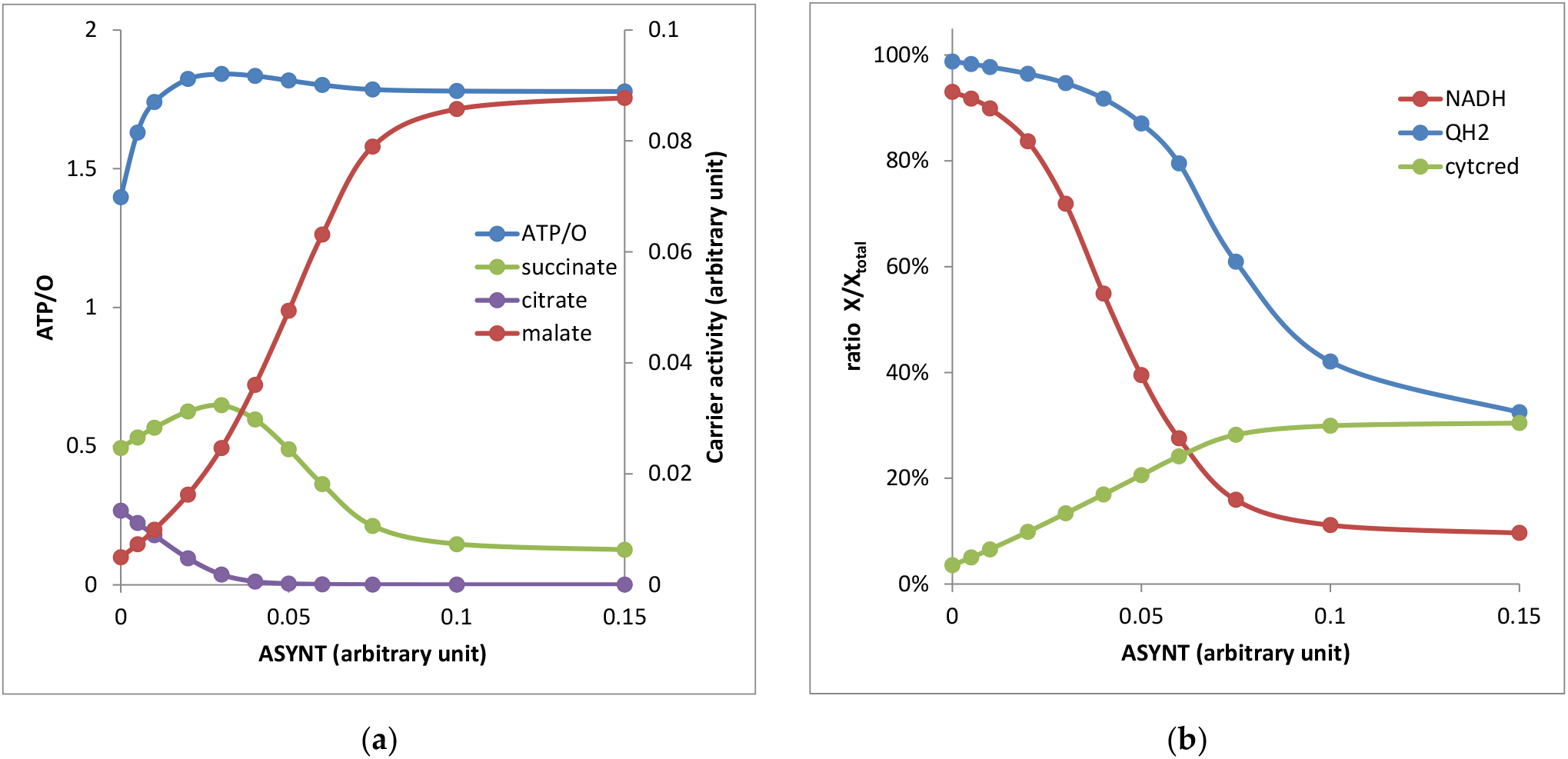
Energy metabolism of isolated yeast mitochondria respiring on AKG as a function of ATP synthase activity. (a) ATP/O ratio (left scale) and output fluxes through Odc1 carrier against AKG input (in arbitrary units, right scale); (b) NADH/NAD_total_ and QH_2_/Q_total_ and reduced cytc/cyt c_total_ ratios. The parameters of the kinetic model are listed in Supplementary Materials S1.

When ASYNT = 0 the respiratory chain activity is low due to the high PMF value leading to high NADH/NAD and QH_2_/Q ratios. The RCII activity is very low so that succinate goes out against AKG. The high NADH level reverses IDH3 activity leading to citrate output against AKG. There is a very low malate output in these conditions. An ATP/O = 1.3 is obtained which fits well with the 1.1 value experimentally observed [20].

An intermediate ASYNT value of 0.04 consumes the PMF which decreases and liberates the respiratory chain which consumes NADH (Figure 2b). The NADH decrease prevents IDH3 reversion and the production of citrate (Figure 2a). The slight decrease of QH_2_ (10%) allows an increase in malate production, but succinate output remains high. An ATP/O = 1.9 is observed in these conditions.

At higher ASYNT activity, above 0.1 (WT), NADH and QH_2_ concentrations become low liberating the RCII activity which consumes a greater part of succinate that is then transformed in fumarate and malate excreted against AKG. The ATP/O is equal to 1.8, a bit lower than the observed value of 2.3. We will discuss this point later.

### 3.3. Decomposition of the steady-states in EFMs (Fig. 3 and Sup. Mat. S2)

Because any set of reactions at steady-state can be expressed as a non-negative combination of the EFMs of the metabolic network, we expressed the different steady-states of figure 2 as a function of the EFMs represented in figure 1. It appeared that only six EFMs are sufficient to describe all the set of fluxes, EFMa, EFMb, EFMd, RFMg, EFMh and EFMi (EFMa means EFM represented in figure 1a etc.). At low ATP synthase activity the set of fluxes is mainly represented by EFMd (output of succinate), EFMg and h (output of succinate and citrate) and EFMi (output of succinate, malate and citrate). As the ATP synthase activity increases, the EFMd, g, h and i are progressively replaced by EFMs a and b then mainly by b (93%) at high ATP synthase activity. These changes are in accordance with the metabolites outputs and the variation of the ATP/O ratio described in figure 2(a).

**Figure 3.**
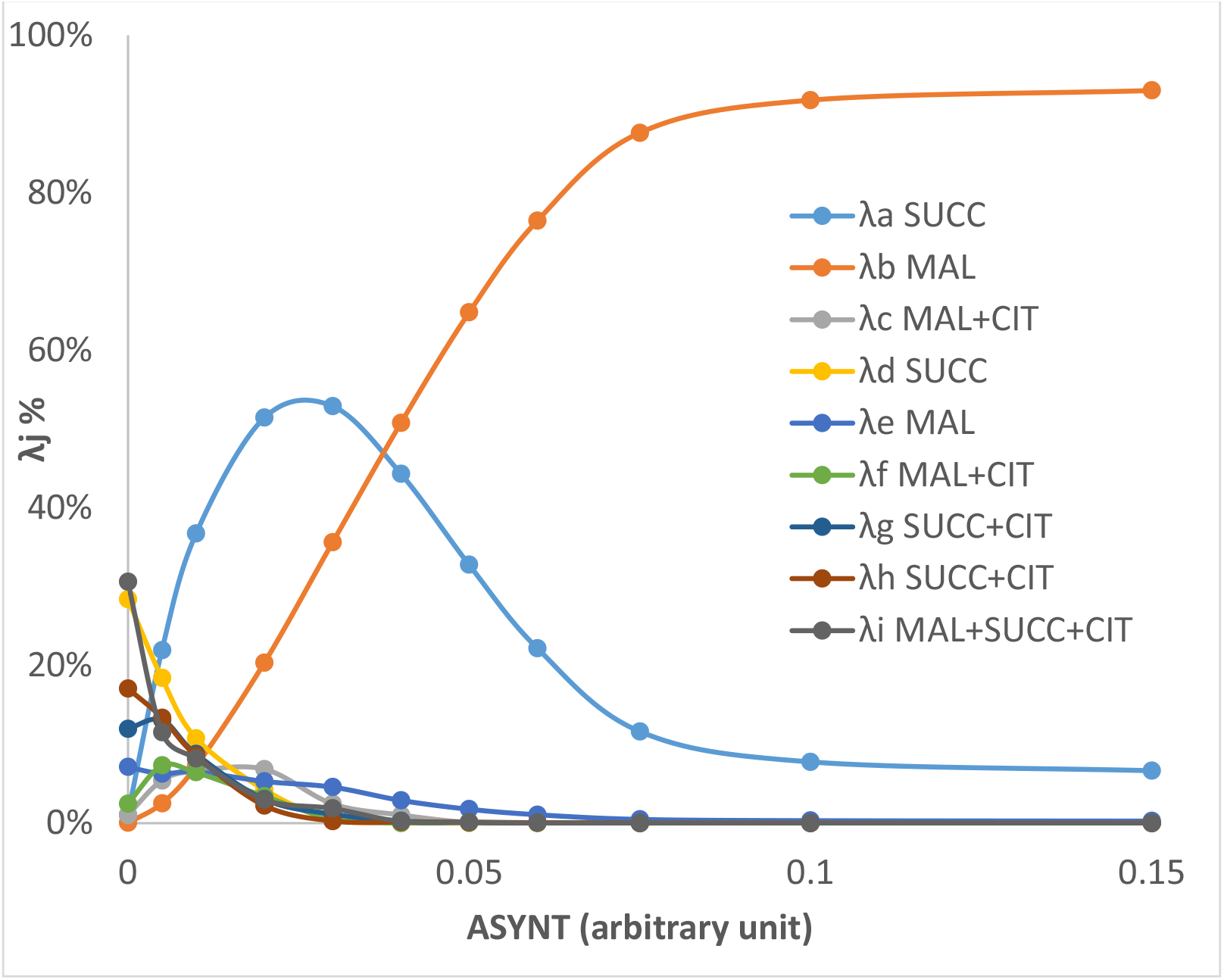
Decomposition of the fluxes as a function of the EFMs of the network involving AKG, ADP and Pi input. (See details in Supp. Mat. S2). The λj values refer to the EFMj represented in the figure 1. For each ASYNT value, the metabolism at steady state is equal to Σλj*EFMj (j=a,b,…i) with Σλj = 1. The metabolite(s) indicated after the λj is (are) the metabolite(s) that goes (go) out.

### 3.4. Overexpression of Odc1p

Curiously, in our model the effect of increasing the activity of Odc1p carrier (x 10) is rather weak (table 1, No Odc1 leak). This could be explained by several factors. First, the growth of the yeast strain NARP + Odc1p is much less than the growth of the wild type (Figure 1 in [8]) suggesting that the increase in ATP synthesis with Odc1p overexpression is far from reaching the WT value ; second, it was shown that in the yeast NARP mutant, the activity of complex IV, the last step of respiratory chain, is decreased by 80%, so that the 0.030 value of ATP synthesis in NARP (Table 1) is probably overestimated. The effect of Odc1p overexpression is not only to increase the expression of the carrier but also to increase the level of the complex IV (40% of the WT activity, table 1 in [8]). This is attributed to the retrograde signaling pathway linking ATP synthase and complex IV biogenesis [8,10,11].

**Table 1.**
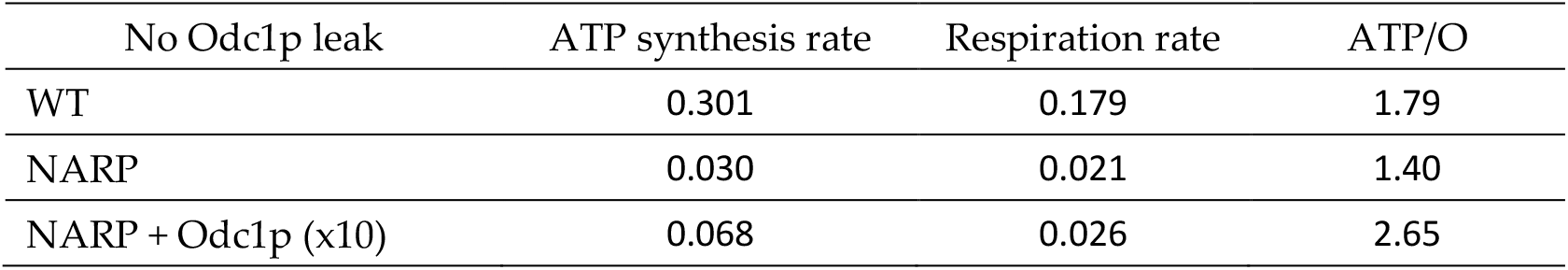
ATP synthesis, respiration and ATP/O in wild type (WT), NARP mutation and NARP + 10 times Odc1p overexpression calculated with Copasi model in arbitrary units.

Another factor could play an important role in the rescue of yeast NARP mutant by Odc1p overexpression : one of the strong constraints of SLP-ATP production is the mandatory reoxidation of the NADH produced by α-ketoglutarate dehydrogenase, the reaction preceding the thiokinase reaction that produces ATP in TCA cycle. Several ways exist in mitochondria to reoxidize NADH [22]. One occurring at low activity of ATP synthase is the reversion of isocitrate dehydrogenase in the presence of high concentration of NADH leading to citrate output as shown in Fig. 2. The other classical one is the activation of the respiratory chain due to the PMF consumption by the ATP synthase. Another way to consume the PMF and NADH, consequently activating the TCA cycle and SLP, is an increase in the membrane leak. In yeast this leak is naturally high leading to a high state 4 and a mild uncoupling between respiration and ATP synthesis (table 1 in [8]). The introduction in our model of a leak accompanying the Odc1 overexpression further increased ATP synthesis rate (table 2).

**Table 2.** ATP synthesis, respiration and ATP/O in wild type (WT), NARP mutation and NARP + 10 times Odc1p overexpression with a leak accompanying Odc1p overexpression. Calculations are performed using the Copasi model in arbitrary units.

| Odc1 leak | ATP synthesis rate | Respiration rate | ATP/O |
| --- | --- | --- | --- |
| WT | 0.300 | 0.169 | 1.77 |
| NARP | 0.031 | 0.034 | 1.11 |
| NARP + Odc1p (x10) | 0.084 | 0.047 | 1.79 |

## 4. Discussion

The introduction of human mitochondrial mutations in yeast is an ingenious strategy to explore curative procedures of mitochondrial diseases. This is how the overexpression of the Odc1p carrier was discovered in yeast [10] with the possible implication of substrate level phosphorylation in the TCA cycle to supply cellular ATP needs.

In order to understand the bases of the rescue of ATP synthase mutants by Odc1p and the metabolic constraints in this metabolic bypass we developed a theoretical model of mitochondrial metabolism with alpha-ketoglutarate (AKG) as respiratory substrate (Supp Mat S1).

### 4.1 EFMs

We first described the Elementary Flux Modes (EFMs) of the metabolic network. EFMs are the minimal pathways in a metabolic network at steady-state. Any set of metabolic fluxes of a metabolic network at steady-state can be expressed as sum of EFMs weighted with non-negative coefficients summing to one.

The description of the EFMs of yeast mitochondrial metabolism showed that, in principle, any TCA segment can be taken from AKG leading to any TCA intermediate output (Fig. 1) with ATP/O ratios ranging from 0.5 to 6 except for EFM of Fig. 1h, for which there is no oxygen consumption. The already reported ATP/O ratio of 2.3 [20] is close to the one of 2.15 measured in EFMs of Fig. 1a and c which correspond in our system to an exit of succinate on the one hand or malate and citrate on the other hand. However, because the metabolic fluxes at steady state can be the combination of several EFMs with different ATP/O values, a value of 2.3 can be achieved in this combination.

### 4.2. Kinetic Model and EFMs decomposition

To get insight in this matter, we introduced in our model the rate equations of the different steps (Supplementary Materials S1) and we studied the steady state of the metabolic network at different ATP synthase activity values (figure 2). We observed a rewiring of mitochondrial metabolism when the amount of ATP synthase is decreased with an ATP/O passing from a value of 1.8-1.9 to a value of 1.3 close to the corresponding experimental values 1.1 reported in [20] in presence of oligomycin, an inhibitor of ATP synthase. However, one may wonder the reason(s) of the greater difference between the theoretical value of 1.8 and the experimental one of 2.3 at high ATP synthase activity. As we noticed earlier, two EFMs present an ATP/O of 2.15 close to the experimental value. One presents an output of citrate which is not observed in our kinetic model at high ATP synthase activity. The other one supposes that all AKG entry occurs in exchange with succinate output. As discussed above, succinate output necessitates that SDH (complex II) is inactive which in our case is obtained at high QH_2_ concentration. Another solution can come from the participation of EFMs with ATP/O = 6. Once more these EFMs involved a citrate output which necessitates high NADH concentration. This is not the case at high ATP synthase concentration in our kinetic model in which the release of metabolites changed from mainly malate and succinate at high ATP synthase activity to succinate plus malate and ultimately, at null ATP synthase activity, to mainly succinate release (plus weaker citrate and malate release). The decomposition of the steady-states into six EFMs (figure 3) confirmed this result, with essentially the EFMb (corresponding to figure 1b and ATP/O = 1.77) and to a less extend EFMa (ATP/O = 2.15) at high ATP synthase activity.

Obvisouly, the conclusions of our model are dependent upon the rate equations of the enzymatic reactions. In order to solve the discrepancy in ATP/O values, experiments are underway in our laboratory to refine the kinetic parameters of our system by measuring the activities of isolated reactions but also the fluxes of metabolites released from the mitochondria with AKG as respiratory substrate.

### 4.3. Odc1p rescue

It has been shown that CCCP, an uncoupler of respiratory chain from ATP synthase activity can also rescue ATP deficiency in yeast [8]. We can thus hypothesize that part of the Odc1p effect could be to increase the membrane leak favoring SLP-ATP not only by increasing AKG feeding part of the TCA cycle producing ATP but also NADH consumption by an activation of the respiratory chain. We tested this hypothesis in our model that showed that increasing the leak in the NARP + Odc1p conditions actually increased SLP-ATP synthesis (table 2). This effect is to be compared with the role of uncoupling proteins in cancer [23,24]. This should also be reconciled with the fact that the overexpression of Odc1 can cure different types of ATP synthase deficiency [8,10,25], probably by a more general mechanism than a simple AKG transport. Work is in progress in our laboratory to study the possible uncoupling properties of the Odc1 carrier.

### 4.4. Implications for NARP syndrome. Human Odc1p counterpart

What is the relevance of this yeast study on human NARP syndrome?

Several reports show that substrate level phosphorylation is able to rescue ATP synthase deficiency or to favor ATP production in cells fed with glutamine, a precursor of AKG [26–28]. However, our theoretical study evidences the constraints in enhancing SLP through a rewiring of energy metabolism, namely the necessity to reoxidize the NADH synthesized by alpha-ketoglutarate dehydrogenase [22,26].

It also evidenced the (perhaps multifactorial) role of the mitochondrial carriers such as Odc1p. Concerning human NARP syndrome, the question is: is there a human homologue of Odc1p? As a matter of fact Odc1p has been identified by sequence analysis of the nuclear genome of *Saccharomyces cerevisiae* [12] and then in human as orthologue [29]. As in yeast, “the human oxodicarboxylate carrier catalyzed the 2-oxoadipate and 2-oxoglutarate by a counter-exchange mechanism”[29]. Nevertheless, contrary to yeast Odc1p, the human carrier does not transport malate and succinate. Thus, the human oxodicarboxylate carrier does not appear as a good candidate to rescue NARP mutations as in yeast. However in their work aiming at rescuing human NARP mutations, Sgarbi et al. [27] were able to increase mitochondrial substrate level phosphorylation and propose a mechanism in which the oxoglutarate carrier exchange 2-oxoglutarate against malate in a medium enriched in 2-oxoglutarate and aspartate. Thus, the human oxoglutarate carrier (OGC) could play the role of the oxodicarboxylate carrier (Odc1) in yeast. The comparison of the fluxes of metabolites released from yeast and mammals mitochondria with AKG as respiratory substrate and different level of ATP synthase activity should help understanding the mitochondrial metabolic pathways taken to rescue ATP synthase deficiencies.

## Supplementary Materials

S1: Rate equations of the theoretical Model with the kinetic parameters; S2: EFMs of the model (figure 1) and decomposition of the steady-sate fluxes on the EFM space (figure 3); S3 sbml file of the copasi model.

## Author Contributions

Conceptualization, A.D., J.-P. M. and S.R.; methodology, J.-P. M. and S.R..; software, J.-P. M. and S.R.; validation, A.D., J.-P. M., S.R. and E.Y.; formal analysis, A.D., J.-P. M., S.R. and E.Y.; writing—original draft, A.D., J.-P. M. and S.R.. All authors have read and agreed to the published version of the manuscript.

## Acknowledgments

This work beneficiated from helpful discussions with Stéphane Duvezin-Caubet, Jean-Paul di Rago, Michel Rigoulet and Déborah Tribouillard-Tanvier

## Conflicts of Interest

The authors declare no conflict of interest.

### Abbreviations

AKG: α-ketoglutarate / 2-oxoglutarate.
ASYNT-ATP: ATP synthesized by the mitochondrial ATP-synthase.
PMF: Proton Motive Force (ΔµH^+^).
SLP: substrate level phosphorylation.
SLP-ATP: ATP synthesized by Succinyl-CoA synthetase (succinate thiokinase).

## References

1. Zhou, A.; Rohou, A.; Schep, D.G.; Bason, J.V.; Montgomery, M.G.; Walker, J.E.; Grigorieff, N.; Rubinstein, J.L. Structure and Conformational States of the Bovine Mitochondrial ATP Synthase by Cryo-EM. Elife 2015, 4, e10180, doi:10.7554/eLife.10180.

2. Dautant, A.; Velours, J.; Giraud, M.-F. Crystal Structure of the Mg·ADP-Inhibited State of the Yeast F1c10-ATP Synthase. J Biol Chem 2010, 285, 29502–29510, doi:10.1074/jbc.M110.124529.

3. Spikes, T.E.; Montgomery, M.G.; Walker, J.E. Structure of the Dimeric ATP Synthase from Bovine Mitochondria. Proc Natl Acad Sci U S A 2020, 117, 23519–23526, doi:10.1073/pnas.2013998117.

4. Guo, H.; Rubinstein, J.L. Cryo-EM of ATP Synthases. Current Opinion in Structural Biology 2018, 52, 71–79, doi:10.1016/j.sbi.2018.08.005.

5. Dautant, A.; Meier, T.; Hahn, A.; Tribouillard-Tanvier, D.; di Rago, J.-P.; Kucharczyk, R. ATP Synthase Diseases of Mitochondrial Genetic Origin. Front Physiol 2018, 9, 329, doi:10.3389/fphys.2018.00329.

6. Su, X.; Dautant, A.; Rak, M.; Godard, F.; Ezkurdia, N.; Bouhier, M.; Bietenhader, M.; Mueller, D.M.; Kucharczyk, R.; di Rago, J.-P.; et al. The Pathogenic m.8993 T > G Mutation in Mitochondrial ATP6 Gene Prevents Proton Release from the Subunit c-Ring Rotor of ATP Synthase. Hum Mol Genet 2021, 30, 381–392, doi:10.1093/hmg/ddab043.

7. Su, X.; Dautant, A.; Godard, F.; Bouhier, M.; Zoladek, T.; Kucharczyk, R.; di Rago, J.-P.; Tribouillard-Tanvier, D. Molecular Basis of the Pathogenic Mechanism Induced by the m.9191T&>C Mutation in Mitochondrial ATP6 Gene. International Journal of Molecular Sciences 2020, 21, 5083, doi:10.3390/ijms21145083.

8. Su, X.; Rak, M.; Tetaud, E.; Godard, F.; Sardin, E.; Bouhier, M.; Gombeau, K.; Caetano-Anollés, D.; Salin, B.; Chen, H.; et al. Deregulating Mitochondrial Metabolite and Ion Transport Has Beneficial Effects in Yeast and Human Cellular Models for NARP Syndrome. Hum Mol Genet 2019, 28, 3792–3804, doi:10.1093/hmg/ddz160.

9. Rak, M.; Tetaud, E.; Duvezin-Caubet, S.; Ezkurdia, N.; Bietenhader, M.; Rytka, J.; di Rago, J.-P. A Yeast Model of the Neurogenic Ataxia Retinitis Pigmentosa (NARP) T8993G Mutation in the Mitochondrial ATP Synthase-6 Gene. J Biol Chem 2007, 282, 34039–34047, doi:10.1074/jbc.M703053200.

10. Schwimmer, C.; Lefebvre-Legendre, L.; Rak, M.; Devin, A.; Slonimski, P.P.; di Rago, J.-P.; Rigoulet, M. Increasing Mitochondrial Substrate-Level Phosphorylation Can Rescue Respiratory Growth of an ATP Synthase-Deficient Yeast. J Biol Chem 2005, 280, 30751–30759, doi:10.1074/jbc.M501831200.

11. Butow, R.A.; Avadhani, N.G. Mitochondrial Signaling: The Retrograde Response. Molecular Cell 2004, 14, 1–15, doi:10.1016/S1097-2765(04)00179-0.

12. Palmieri, L.; Agrimi, G.; Runswick, M.J.; Fearnley, I.M.; Palmieri, F.; Walker, J.E. Identification in Saccharomyces Cerevisiae of Two Isoforms of a Novel Mitochondrial Transporter for 2-Oxoadipate and 2-Oxoglutarate. J Biol Chem 2001, 276, 1916–1922, doi:10.1074/jbc.M004332200.

13. Lebiedzinska, M.; Karkucinska-Wieckowska, A.; Wojtala, A.; Suski, J.M.; Szabadkai, G.; Wilczynski, G.; Wlodarczyk, J.; Diogo, C.V.; Oliveira, P.J.; Tauber, J.; et al. Disrupted ATP Synthase Activity and Mitochondrial Hyperpolarisation-Dependent Oxidative Stress Is Associated with P66Shc Phosphorylation in Fibroblasts of NARP Patients. Int J Biochem Cell Biol 2013, 45, 141–150, doi:10.1016/j.biocel.2012.07.020.

14. Hoops, S.; Sahle, S.; Gauges, R.; Lee, C.; Pahle, J.; Simus, N.; Singhal, M.; Xu, L.; Mendes, P.; Kummer, U. COPASI--a COmplex PAthway SImulator. Bioinformatics 2006, 22, 3067–3074, doi:10.1093/bioinformatics/btl485.

15. Bohnensack, R. Control of Energy Transformation of Mitochondria. Analysis by a Quantitative Model. Biochim Biophys Acta 1981, 634, 203–218, doi:10.1016/0005-2728(81)90139-0.

16. Mitchell, P. Vectorial Chemistry and the Molecular Mechanics of Chemiosmotic Coupling: Power Transmission by Proticity. Biochemical Society Transactions 1976, 4, 399–430, doi:10.1042/bst0040399.

17. Toth, A.; Meyrat, A.; Stoldt, S.; Santiago, R.; Wenzel, D.; Jakobs, S.; Ballmoos, C. von; Ott, M.Kinetic Coupling of the Respiratory Chain with ATP Synthase, but Not Proton Gradients, Drives ATP Production in Cristae Membranes. PNAS 2020, 117, 2412–2421, doi:10.1073/pnas.1917968117.

18. Schuster S; Hilgetag, C. On Elementary Flux Modes in Biochemical Reaction Systems at Steady State. J. Biol. Syst. 1994, 165–182.

19. Pfeiffer, T.; Sánchez-Valdenebro, I.; Nuño, J.C.; Montero, F.; Schuster, S. METATOOL: For Studying Metabolic Networks. Bioinformatics 1999, 15, 251–257, doi:10.1093/bioinformatics/15.3.251.

20. Rigoulet, M.; Velours, J.; Guerin, B. Substrate-Level Phosphorylation in Isolated Yeast Mitochondria. Eur J Biochem 1985, 153, 601–607, doi:10.1111/j.1432-1033.1985.tb09343.x.

21. Schuster, S.; Dandekar, T.; Fell, D.A.; Schuster, S.; Dandekar, T.; Fell, D.A.; Schuster, S.; Dandekar, T.; Fell, D.A.; Schuster, S.; et al. Detection of Elementary Flux Modes in Biochemical Networks: A Promising Tool for Pathway Analysis and Metabolic Engineering. Trends in Biotechnology 1999, 17, 53–60, doi:10.1016/S0167-7799(98)01290-6.

22. Chinopoulos, C. Acute Sources of Mitochondrial NAD+ during Respiratory Chain Dysfunction. Experimental Neurology 2020, 327, 113218, doi:10.1016/j.expneurol.2020.113218.

23. Valle, A.; Oliver, J.; Roca, P. Role of Uncoupling Proteins in Cancer. Cancers (Basel) 2010, 2, 567–591, doi:10.3390/cancers2020567.

24. Samudio, I.; Fiegl, M.; Andreeff, M. Mitochondrial Uncoupling and the Warburg Effect: Molecular Basis for the Reprogramming of Cancer Cell Metabolism. Cancer Res 2009, 69, 2163–2166, doi:10.1158/0008-5472.CAN-08-3722.

25. de Taffin de Tilques, M.; Tribouillard-Tanvier, D.; Tétaud, E.; Testet, E.; di Rago, J.-P.; Lasserre, J.-P. Overexpression of Mitochondrial Oxodicarboxylate Carrier (ODC1) Preserves Oxidative Phosphorylation in a Yeast Model of Barth Syndrome. Dis Model Mech 2017, 10, 439–450, doi:10.1242/dmm.027540.

26. Chinopoulos, C.; Seyfried, T.N. Mitochondrial Substrate-Level Phosphorylation as Energy Source for Glioblastoma: Review and Hypothesis. ASN Neuro 2018, 10, 1759091418818261, doi:10.1177/1759091418818261.

27. Sgarbi, G.; Casalena, G.A.; Baracca, A.; Lenaz, G.; DiMauro, S.; Solaini, G. Human NARP Mitochondrial Mutation Metabolism Corrected with Alpha-Ketoglutarate/Aspartate: A Potential New Therapy. Arch Neurol 2009, 66, 951–957, doi:10.1001/archneurol.2009.134.

28. Chen, Q.; Kirk, K.; Shurubor, Y.I.; Zhao, D.; Arreguin, A.J.; Shahi, I.; Valsecchi, F.; Primiano, G.; Calder, E.L.; Carelli, V.; et al. Rewiring of Glutamine Metabolism Is a Bioenergetic Adaptation of Human Cells with Mitochondrial DNA Mutations. Cell Metab 2018, 27, 1007-1025.e5, doi:10.1016/j.cmet.2018.03.002.

29. Fiermonte, G.; Dolce, V.; Palmieri, L.; Ventura, M.; Runswick, M.J.; Palmieri, F.; Walker, J.E. Identification of the Human Mitochondrial Oxodicarboxylate Carrier. Bacterial Expression, Reconstitution, Functional Characterization, Tissue Distribution, and Chromosomal Location. J Biol Chem 2001, 276, 8225–8230, doi:10.1074/jbc.M009607200.

