## Supplementary material for "A theoretical model of mitochondrial ATP synthase deficiencies. The role of mitochondrial carriers": ARTICLE MITOY Rev Supp Mat S1-2.pdf

**Supplementary Material S1:**

**Copasi model Ransac-Mazat 2021 MITOYeast_AKG_12_v14.cps**

**INTRODUCTION.**

According to [1], we define as global quantities in Copasi, 'fmu' which determines the part of $\Delta\psi$ (DpSI) in $\Delta\mu H^+$ (PMF). Here we take for simplicity: fmu = 0.8 and thus DPSI = fmu*PMF (Volts) and DPH = (1-fmu)*PMF/Z (in pH units).
We also define Z = ln(10)*RT/F ≈ 0.06 Volts at 30°C.
All stoichiometric coefficients are integer, which is necessary for the calculation of the EFMs (Elementary Flux Modes).

**α-KETOGLUTARATE DEHYDROGENASE (AKGDH): AKGm + NADm + CoAm -> SucCoAm + NADHm + CO2**

It is supposed irreversible (ΔG°= -27.2 ± 7.7 kJ/mol => K'eq = 5.9 × $10^4$) and inhibited by its products NADH and SucCoA. We propose an irreversible mass action law for the substrates, modulated by competitive inhibition terms for NADHm and SucCoAm with KAKGdh_NADH = 0.03 mM and KAKGdh_SucCoA = 10 µM taken from the literature [2–4]:

$$VM_{AKGDH}.AKGm.NADm.CoAm.\frac{1}{1+\frac{NADHm}{KAKGdh\_NADH}}.\frac{1}{1+\frac{SucCoAm}{KAKGdh\_SucCoA}}$$

or linearly written:
VM_AKGDH*AKGm*NADm*CoAm*(1/(1+NADHm/KAKGdh_NADH))*(1/(1+SucCoAm/KAKGdh_SucCoA))

**ANT: 5 * ADPc + 5 * ATPm + 4 * PMF = 5 * ADPm + 5 * ATPc**

We take the reversible mass action law with the intervention of DPSI = fmu*PMF

vANT = VM_ANT*(ADPc*ATPm-ADPm*ATPc*10^(-fmu*PMF))

**ASYNT**

**ATP SYNTHASE (ASYNT) : 3 * ADPm + 3 * Pim + 10 * PMF = 3 * ATPm**
(stoichiometry of yeast ATP synthase: 10 $H^+$ per 3 ATP).
We took the expression of Bohnensack [1]:

$$VM_{ASYNT}.\left(1-\frac{ATPm}{ADPm.Pim}.Phi_p.10^{nA.\frac{PMFa-PMF}{Z}}\right)$$

where nA is the number of protons translocated per ATP synthesised (3.3 in the case of yeast), Phi_p takes into account the DPH in the phosphate ions equilibrium (see[1]) and PMFa is the standard phosphorylation potential (0.150 V) [1].

**ATPASE : ATPc -> ADPc + Pic**

We take a Henri-Michaelis-Menten equation [5] with the ATPc as substrate:

$$VM\_ATPASE.\frac{\frac{ATPc}{KATPASE\_ATP}}{1+\frac{ATPc}{KATPASE\_ATP}}$$

**CITRATE SYNTHASE (CS) : AcCoAm + 0AAm = CITm + CoAm**

We consider the reaction as reversible and take the rate expression of Cornish-Bowden and Hofmeyr [6] with or without the inhibition by ATP (the inhibition by CoA is supposed to be included in the reverse rate)

$$V=\frac{\frac{VMCS.[OAAm]\times[ACoAm]}{KCS\_OAA\times KCS\_ACOA}\left(1-\frac{[CoA]\times[CITm]}{Kq\_CS\times[ACoAm]\times[OAAm]}\right)}{\left[1+\frac{[OAA]}{KCS\_OAA}+\frac{[AcCoA]}{KCS\_ACOA}\left[1+\frac{ATP}{KiCS\_ATP}\right]\right].\left[1+\frac{[CITm]}{KCS\_CIT}+\frac{[CoA]}{KCS\_COA}.\right]}$$

with the consensus values (in mM) : KCS_OAA = 0.005 ; KCS_ACOA = 0.01 ; KQ_CS = 2.10^6 (Veech JBC 1973) ; KiCS_ATP = 0.3 ; KCS_CIT = 3 ; KCS_COA = 0.04.

**FUMARASE: FUMm = MALm**

Reversible Henri-Michaelis-Menten equation, which takes into account the equilibrium constant KQ_FUMASE = 4.4 in order to satisfy Haldane relationship:

(KFUMASE_FUM = 0.004 mM et KFUMASE_MAL = 0.009 mM)

$$VM\_FUM.\frac{\left(\frac{FUMm}{KFUMASE\_FUM}-\frac{MALm}{KQ\_FUMASE.KFUMASE\_FUM}\right)}{1+\frac{FUMm}{KFUMASE\_FUM}+\frac{MALm}{KFUMASE\_MAL}}$$

**IDH3: ACONITASE-ISOCITRATE DEHYDROGENASE 3**

Aconitase and isocitrate dehydrogenase are joined together in the reaction:

**CITm + NADm = AKGm + NADHm + CO2**

and the rate equation:

$$V=VM\_IDH3\frac{\frac{.[CITm]\times[NADm]}{KI3\_CIT\times KI3\_NAD}\left(1-\frac{[AKGm]\times[NADHm]}{Kq\_IDH3\times[CITm]\times[NADm]}\right)}{\left[1+\frac{[CITm]}{KI3\_CIT}+\frac{[AKGm]}{KI3\_NAD}\right]\times\left[1+\frac{[NADm]}{KI3\_NAD}+\frac{[NADHm]}{KI3\_NADH}.\right]}$$

**LEAK: PMF -> PMFm**

We take the equation:

$$\frac{VM\_Leak}{1+\frac{KLEAK}{10^{\frac{PMF}{Z}}}}$$

**MDH MALATE DEHYDROGENASE (Mitochondrial: MDH2): MALm + NADm = OAAm + NADHm**

Mass Action reversible with Keq = 0.0002

**MALIC ENZYME (ME2): Malate + NAD(P)+ ⇌ pyruvate + CO2 + NAD(P)H.**

Mass action reversible with Keq (NAD ou NADPH) = 7.6 $10^{-3}$

**NDI: NADHm + Q -> NADm + QH2**

In yeast, NDI is the internal dehydrogenase, which takes the place of complex I in mammal's mitochondria. Because it does not excrete protons, its equilibrium constants $K_{eq}$= $4.10^{13}$ can be calculated from the difference in the Q/QH2 and NAD/NADH midpoint potentials (+0.085 V and -0.32 V respectively).

Because the $K_{eq}$ value, it is supposed irreversible, we take k1 = 1 $s^{-1}$ in a irreversible mass action law.

**PYRUVATE DEHYDROGENASE (PDH) : $NAD^+$ + CoA + Pyruvate -> NADH + CO2 + Acetyl-CoA**
**NADm + CoAm + PYRm -> NADHm + CO2 + ACoAm**
We take the equation given in [7]. This reaction with a complex mechanism is taken as irreversible despite its ΔG° = -35.3 ± 6.4 kJ/mol ($K'_{eq}$ = 1.6 × $10^6$ ). We take into account an inhibition by the products NADH and acetyl-CoA (ACoA) (which is more or less equivalent to a reversible mechanism):

$$\frac{\boldsymbol{v}}{\boldsymbol{V_{max}}} = \boldsymbol{VMPDH} * \frac{\boldsymbol{CoAm}}{\boldsymbol{KPDHCoAm} + \boldsymbol{CoAm}} * \frac{\boldsymbol{PYRm}}{\boldsymbol{KPDHPYR} + \boldsymbol{PYRm}} * \frac{\boldsymbol{NADm}}{\boldsymbol{KPDHNAD} * \left(\boldsymbol{1} + \frac{\boldsymbol{NADHm}}{\boldsymbol{KNADH}}\right) + \boldsymbol{NADm}} * \frac{\boldsymbol{KiACoA}}{\boldsymbol{KiACoA} + \boldsymbol{ACoAm}}$$

**Linearly written, the equation reads:**
VMPDH*(CoAm/(KPDHCoAm+CoAm))*(PYRm/(KPDHPYR+PYRm))*(NADm/(KPDHNAD*(1+NADHm/KiNADH)+NADm))*(KiACoA/(KiACoA+ACoAm))

**Pi TRANSPORT (Pit): 5 * Pic + PMF = 5 * Pim**
The electroneutral transport of Pi with a $H^+$ (or against $OH^-$) is very fast so that the phosphate distribution is supposed to be in equilibrium (taking into account the ΔpH).

According to[1] we take:

$$V\_Pit = VM\_Pit.\left(1 - \frac{Pim}{Pic}.10^{-DPH}\right) = VM\_Pit.\left(1 - \frac{Pim}{Pic}.10^{-\frac{0.2.PMF}{Z}}\right)$$

or linearly written:

V_Pit = VM_Pit*(1-Pim/Pic*10^(-0.2*PMF/Z))

**RCII (Respiratory complex II) or SDH (Succinate dehydrogenase): SUCCm + Q = FUMm + QH2; OAAm**

Keq = 70 is calculated from the difference in the midpoint potentials of the pairs Fumarate/Succinate (E1 = +0.03 V) and Q/QH2 (E2 = + 0.085 V).

We adopt the rate equation of Cornish-Bowden and Hofmeyr [6] with a competitive inhibition of oxaloacetate with succinate and fumarate:

$$V = VM\_SDH \times \frac{\frac{[SUCCm]\times[Q]}{KSDH\_SUCC\times\left[1+\frac{[OAAm]}{KiSDH\_OAA}\right]\times KSDH\_Q}\left(1-\frac{[QH2][FUMm]}{KQ\_SDH\times[SUCCm]\times[Q]}\right)}{\left[1+\frac{[SUCCm]}{KSDH\_SUCC\times\left[1+\frac{[OAAm]}{KiSDH\_OAA}\right]}+\frac{[FUMm]}{KSDH\_FUM\times\left[1+\frac{[OAAm]}{KiSDH\_OAA}\right]}\right]\times\left[1+\frac{[Q]}{KSDH\_Q}+\frac{[QH2]}{KSDH\_QH2}.\right]}$$

With the consensus values (for mammalian ): KSDH_SUCC = 85 µM, $K_{SDH\_FUM}$ = 100 µM, $K_{SDH\_Q}$ = 1.5 µM (Grivennikova et al., [8]) and $K_{SDH\_QH2}$ = 1.5 µM (arbitrarily equal to $K_M^Q$ = 1.5 µM in absence of known determination). KiSDH_OAA_ = 0.07 µM (0.01< <0.4)

Note that oxaloacetate (OAAm) is not a metabolite of our model

**RCIII or bc1 complex: 5 * QH2 + 10 * Cox = 5 * Q + 10 * Cred + 12 * PMF** (4 protons (4x0.2 = 0.8 PMF) and 2 charges (2x0.8=1.6))*5.

We follow the model of Bernard Korzeniewski [9] which was also taken over by Daniel A. Beard [8] of a linear dependence of the rate upon the difference of potential near equilibrium:

VCIII = VM_CIII * ΔECIII.

ΔECIII (for two electrons) decomposes into a $\Delta ECIII_{Chem}$ =2(EC – EQ) which accounts for the chemical part of the reaction and –2.4 * PMF (Volts) corresponding to the protons and charges transported on either side of the inner mitochondrial membrane

ΔECIII = 2(EC – EQ) – 2.4.PMF = 2 ([$EC_0$ + RT/F *ln(Cox/Cred)] – [$EQ_0$ + RT/2F*ln(Q/QH2)]) – 2.4.PMF

With $EC_0$ (standard potential of Cred/Cox) = + 0.25 V and $EQ_0$ (standard potential of Q/QH2) = +0.085 V and with RT/F. ln(10) = 0.06 V at 30°C, we obtain:

ΔECIII = 2[0.25 + 0.06.$\log_{10}$(Cox/Cred) – (0.085+ 0.03. $\log_{10}$(Q/QH2)] – 2.4.PMF

Linearly written the equation reads:
VM_CIII*(0.50+0.12*$log_{10}$(Cox/Cred))-(0.17+0.06*$log_{10}$(Q/QH2))-2.4*PMF)

**RCIV or Cytochrome c oxidase. 20 * Cred + 5 * O2 -> 20 * Cox + 10 * H2O + 36 * PMF**

We take an equation close to that of Korzeniewski ((6)):

$$\frac{VM_{CIV} * Cred}{\frac{KO}{O2} + 1}$$

**SUCCINATE THIOKINASE or SUCCINYL-COA SYNTHETASE (SUCTHIOK)**: **SUCCoAm + ADPm + Pim = SUCCm + ATPm + CoAm**

We consider this equation as reversible with KQ_STK = 1.3 integrated in the mass action law equation:

VM_STK*(SUCCoAm*ADPm*Pim -(SUCCm*ATPm*CoAm)/(KQ_STK))

**ODC1 or ODC2 (Oxo-dicarboxylate carrier)**

**T2**: AKGc + MALm -> AKGm + MALc . (ODC)

Mass action irreversible

**T6:** 5 * PYRm -> 5 * PYRc + PMF

Mass action irreversible

**T22**: AKGc + SUCCm -> AKGm + SUCCc . (ODC)

Mass action irreversible

**T23**: AKGc + CITm -> AKGm + CITc (ODC)

Mass action reversible

T2, T22 and T23 are taken irreversible because MAL, SUCC ant CIT are not present outside mitochondria in our model.

As an indication, Odc1 exchanges the following internal metabolites against AKG ext (with the corresponding rate in mmol.min/mg prot): AKG (112), Malate (56), Succinate (19). [11]

DIC (Dicarboxylate carrier)

**T31:** MALm + Pic = MALc + Pim .

Mass action irreversible

**T33:** SUCCc + Pim = SUCCm + Pic .

Mass action irreversible

**Parameters of the model:**

| | | | |
|---|---|---|---|
| P | 0 | (RCIII).VM_CIII mmol/(l*s) | 2 |
| Z | 0.06 | (RCIV).VM_CIV 1/s | 0.05 |
| DGP | 0.380956 | (RCIV).KO mmol/l | 0.001 |
| nA | 3.333 | (LEAK).VM_Leak mmol/(l*s) | 0.15 |
| DG_SN | 0.246103 | (LEAK).fleakodc1 ? | 0 |
| Gamma | 12639.2 | (LEAK).fleakasynt ? | 0 |
| fmu | 0.8 | (LEAK).KLEAK 1 | 10000 |
| DPSI | 0.150509 | (LEAK).VMASYNT ? | 0 |
| DPH | 0.627122 | (LEAK).VMODC1 ? | 0.0467299 |
| Ac | 5.00E-13 | (LEAK).Z mmol/l | 0.06 |
| Am | 0.0299472 | (T2).k1 l/(mmol*s) | 1 |
| fPSI | 0.8 | (NDI).k1 l/(mmol*s) | 1 |
| Nm | 0.13973 | (RCII).VM_SDH mmol/(l*s) | 1 |
| fp | 0.2 | (RCII).KSDH_SUCC mmol/l | 0.085 |
| Phi_p | 3.59009 | (RCII).KiSDH_OAA mmol/l | 0.07 |
| VMASYNT | 0 | (RCII).KSDH_Q mmol/l | 0.0015 |
| VMODC1 | 1.96 | (RCII).KQ_SDH 1 | 70 |
| Kslip | 1.00E-07 | (RCII).KSDH_FUM mmol/l | 0.1 |
| Pslip | 0 | (RCII).KSDH_QH2 mmol/l | 0.0015 |
| KO2 | 0.001 | (T22).k1 l/(mmol*s) | 0.4 |
| VM_CIV | 0.05 | (T31).k1 l/(mmol*s) | 0 |
| VMODC1_bis | 0.0467299 | (T33).k1 l/(mmol*s) | 0 |
| Koligo | 0.001 | (CS).VM_CS mmol/(l*s) | 11.5 |
| k1succ | 0.4 | (CS).KCS_OAA mmol/l | 0.005 |
| k1cit | 0.56 | (CS).KCS_ACOA mmol/l | 0.01 |
| fsucc | 0.4 | (CS).KQ_CS 1 | 2.00E+06 |
| fcit | 0.56 | (CS).KiCS_ATP mmol/l | 0.3 |
| quantity | 0 | (CS).KCS_CIT mmol/l | 3 |
| ATP/O | 0.346216 | (CS).KCS_COA mmol/l | 0.04 |
| (AKGDH).VM_AKGDH lÂ²/(mmolÂ²*s) | 340 | (IDH3).VM_IDH3 mmol/(l*s) | 5.8 |
| (AKGDH).KAKGdh_NADH mmol/l | 0.03 | (IDH3).KI3_CIT mmol/l | 3.2 |
| (AKGDH).KAKGdh_SucCoA mmol/l | 0.01 | (IDH3).KI3_NAD mmol/l | 0.2 |
| (SUCTHIOK).VM_STK lÂ²/(mmolÂ²*s) | 6.97225 | (IDH3).KQ_IDH3 1 | 7 |
| (SUCTHIOK).KQ_STK 1 | 1.3 | (IDH3).KI3_AKG mmol/l | 9 |
| (FUMARASE).VM_FUM mmol/(l*s) | 12.5478 | (IDH3).KI3_NADH mmol/l | 0.7 |
| (FUMARASE).KFUMASE_FUM mmol/l | 0.004 | (MDH2).k1 l/(mmol*s) | 20 |
| (FUMARASE).KQ_FUMASE 1 | 4 | (MDH2).k2 l/(mmol*s) | 100000 |
| (FUMARASE).KFUMASE_MAL mmol/l | 0.009 | (ME2).k1 l/(mmol*s) | 0 |
| (ASYNT).VM_ASYNT mmol/(l*s) | 0 | (ME2).k2 lÂ²/(mmolÂ²*s) | 0 |
| (ASYNT).PMFa mmol/l | 0.15 | (PDH).VMPDH mmol/(l*s) | 7 |
| (ASYNT).Koligo mmol/l | 0.001 | (PDH).KPDHPYR mmol/l | 0.025 |
| (ASYNT).Phi_p mmol/l | 3.59009 | (PDH).KPDHNAD mmol/l | 0.05 |
| (ASYNT).Z ? | 0.06 | (PDH).KiNADH mmol/l | 0.036 |
| (ASYNT).nA ? | 3.333 | (PDH).KPDHCoAm mmol/l | 0.013 |
| (ANT).VM_ANT l/(mmol*s) | 0.2 | (PDH).KiACoA mmol/l | 0.035 |

| | | | |
|---|---|---|---|
| (ANT).fmu l/mmol | 0.8 | (T6).k1 l^4/(mmol^4*s) | 1 |
| (Pit).VM_Pit mmol/(l*s) | 200 | (T24).k1 1/s | 100 |
| (Pit).Z mmol/l | 0.06 | (T24).k2 1/s | 100 |
| (ATPASE).VM_ATPASE mmol/(l*s) | 0.05 | (T23).k1 l/(mmol*s) | 0.56 |
| (ATPASE).KATPASE_ATP mmol/l | 3.57734 | | |

**References**


1. Bohnensack, R. Control of Energy Transformation of Mitochondria. Analysis by a Quantitative Model. *Biochim Biophys Acta* **1981**, *634*, 203–218, doi:10.1016/0005-2728(81)90139-0.
2. Yang, L.; Garcia Canaveras, J.C.; Chen, Z.; Wang, L.; Liang, L.; Jang, C.; Mayr, J.A.; Zhang, Z.; Ghergurovich, J.M.; Zhan, L.; et al. Serine Catabolism Feeds NADH When Respiration Is Impaired. *Cell Metab.* **2020**, *31*, 809-821.e6, doi:10.1016/j.cmet.2020.02.017.
3. Hamada, M.; Koike, K.; Nakaula, Y.; Hiraoka, T.; Koike, M. A Kinetic Study of the Alpha-Keto Acid Dehydrogenase Complexes from Pig Heart Mitochondria. *J Biochem* **1975**, *77*, 1047–1056, doi:10.1093/oxfordjournals.jbchem.a130805.
4. Smith, C.M.; Bryla, J.; Williamson, J.R. Regulation of Mitochondrial Alpha-Ketoglutarate Metabolism by Product Inhibition at Alpha-Ketoglutarate Dehydrogenase. *J Biol Chem* **1974**, *249*, 1497–1505.
5. Henri Victor Theorie Generale de l’action de Quelques Diastases. *C R Hebd Seances Acad Sci 135, 916–919* 1902.
6. Cornish-Bowden A; Hofmeyr J. Enzymes in context: Kinetic characterization of enzymes for systems biology. The Biochemist April 2005: 11-14. Available online: https://www.qwant.com/ (accessed on 27 June 2021).
7. König, M.; Bulik, S.; Holzhütter, H.-G. Quantifying the Contribution of the Liver to Glucose Homeostasis: A Detailed Kinetic Model of Human Hepatic Glucose Metabolism. *PLoS Comput Biol* **2012**, *8*, e1002577, doi:10.1371/journal.pcbi.1002577.
8. Grivennikova, V.G.; Gavrikova, E.V.; Timoshin, A.A.; Vinogradov, A.D. Fumarate Reductase Activity of Bovine Heart Succinate-Ubiquinone Reductase. New Assay System and Overall Properties of the Reaction. *Biochim Biophys Acta* **1993**, *1140*, 282–292, doi:10.1016/0005-2728(93)90067-p.
9. Korzeniewski, B.; Zoladz, J.A. A Model of Oxidative Phosphorylation in Mammalian Skeletal Muscle. *Biophysical Chemistry* **2001**, *92*, 17–34, doi:10.1016/S0301-4622(01)00184-3.
10. Beard, D.A. A Biophysical Model of the Mitochondrial Respiratory System and Oxidative Phosphorylation. *PLoS Comput Biol* **2005**, *1*, e36, doi:10.1371/journal.pcbi.0010036.
11. Palmieri, L.; Agrimi, G.; Runswick, M.J.; Fearnley, I.M.; Palmieri, F.; Walker, J.E. Identification in Saccharomyces Cerevisiae of Two Isoforms of a Novel Mitochondrial Transporter for 2-Oxoadipate and 2-Oxoglutarate. *J Biol Chem* **2001**, *276*, 1916–1922, doi:10.1074/jbc.M004332200.
